# Spatial Patterns of White Matter Hyperintensities and Cognitive Function in Older Adults: A Community-Based Study

**DOI:** 10.64898/2026.10.14.759131

**Authors:** Xingyu Chen, Kei Nishimaki, Shuhei Tomoshige, Jill S. Chotiyanonta, Yuto Uchida, Bruce A. Wasserman, Rebecca F. Gottesman, Eliseo Guallar, B. Gwen Windham, Kenichi Oishi

## Abstract

**Objectives:** White matter hyperintensities (WMH) are associated with cognitive decline in older adults. We examined whether anatomically defined spatial WMH patterns are associated with cognitive performance and subsequent decline in a large community-based cohort with approximately six years of follow-up.

**Methods:** We analyzed 1,667 participants (mean age 76 years) from the Atherosclerosis Risk in Communities Neurocognitive Study. WMH were segmented from FLAIR images and mapped to 112 white matter regions using deep-learning-based parcellation. Principal component analysis identified dominant WMH distribution patterns; candidate components were selected by Horn’s parallel analysis and retained if stable across resampled split samples. Linear mixed-effects models examined associations between WMH patterns and cognitive performance at baseline and follow-up, adjusting for demographic, genomic, and vascular risk factors. Robustness was assessed using analyses accounting for total WMH burden, study center, and selective attrition.

**Results:** Of 14 candidate principal components (PCs) identified by parallel analysis, eight met the resampling stability criterion and were retained, together explaining 42% of the variance. Three components were associated with cross-sectional or future cognition. PC3, characterized by deep white matter burden in the corpus callosum and anterior corona radiata with relative preservation of superficial white matter, showed no cross-sectional association at baseline but was associated with accelerated decline across multiple domains.

**Interpretation:** A deep spatial pattern of WMH loading on the corpus callosum and anterior corona radiata was associated with subsequent cognitive decline in the absence of a cross-sectional association at baseline, independent of total lesion burden.

## Introduction

White matter hyperintensities (WMH) are abnormal, hyperintense regions observed on T2-weighted and fluid-attenuated inversion recovery (FLAIR) magnetic resonance imaging (MRI) scans. These lesions reflect white matter injury, including demyelination and axonal loss^1, 2^, and are frequently seen in older adults and individuals with vascular risk factors such as hypertension, diabetes, and cerebrovascular disease^3^. Greater WMH burden is consistently associated with poorer cognition and increased risk of cognitive decline, mild cognitive impairment, and dementia^4–6^. Accordingly, total WMH volume has become the predominant imaging measure in studies of cognitive aging^7, 8^.

Accumulating evidence, however, indicates that WMH location may provide information beyond global lesion burden. Diffusion MRI linked WMH affecting major white matter pathways to impaired network efficiency and cognitive slowing^9^. Regional approaches have similarly associated lobar and radial WMH distributions with episodic memory and processing speed^10^. More recently, data-driven studies have characterized WMH topography without predefined regional hypotheses. Phuah et al. used voxel-based spectral clustering to identify reproducible WMH spatial signatures associated with distinct small-vessel disease etiologies^11^, while Rotblatt et al. identified regional WMH profiles associated with different longitudinal cognitive trajectories^12^. Together, these studies establish that WMH topography contains potentially meaningful information beyond total lesion burden.

Several limitations remain. First, the spatial units employed lack detailed anatomical definition. Geometric partitions such as the bullseye method rely on simplified radial and angular divisions^10^, while lobar WMH volumes aggregate structurally and functionally heterogeneous tracts^12^. Data-driven clustering approaches also spans across regions and are not fully defined with reference to known anatomy^11^. Second, prior approaches either categorize WMH patterns into coarse territories, or assign participants into subgroups, which could reduce spatial information and obscure within-group variation. Third, the available evidence, derived from memory-clinic or research-center cohorts, is either cross-sectional or limited to approximately two years of follow-up^12^. The extent to which anatomically fine-grained, continuously measured WMH patterns are reproducible in large community-based populations and relate to cognitive trajectories over longer follow-up remains less well established.

To address these limitations, we analyzed the spatial distribution of WMH in a community-based cohort^13^ with approximately six years of cognitive follow-up. We segmented WMH from FLAIR images and parcellated the brain using a deep learning-based algorithm^14^ into 112 anatomically defined regions. Rather than assigning participants to discrete subgroups, we quantified region-wise WMH prevalence and applied principal component analysis to derive continuous spatial components, and examined their relationship to domain-specific cognitive performance at multiple timepoints while adjusting for relevant covariates. Using this anatomically grounded and continuously measured approach, we examined how the spatial distribution of WMH relates to domain-specific cognitive trajectories over time.

## Methods

### Participants and design

The data used in this study were obtained from the Atherosclerosis Risk in Communities (ARIC) Study, available subject to ARIC data use agreements and approval procedures. No additional IRB approval is required. ARIC is a prospective epidemiological cohort study designed to investigate atherosclerosis from midlife. Participants from 4 different communities had their longitudinal data collected through a series of visits, starting from enrollment between 1987 and 1989 when participants were 45 to 64 years of age ^13^. Then, from 2011 to 2013, a comprehensive dementia surveillance study, the Atherosclerosis Risk in Communities Neurocognitive Study (ARIC-NCS), was initiated at Visit 5 to more comprehensively assess cognitive impairment. Among the 6,538 participants who attended the visit, 1,980 participants were selected for the MRI examination. The selection was based on a probability sampling mechanism, with oversampling that included all participants who had cognitive decline or an MRI examination before Visit 5^15^. The MRI data collection in ARIC adheres to protocols that ensure consistency and reliability across study sites, and the methodologies are standardized with quality control measures^16^. Follow-up neurocognitive assessments were conducted during Visit 6 (2016-2017), 7 (2018-2019), or both.

Brain images were obtained using 3T MRI scanners with a standard set of sequences that included a T1-weighted 3D volumetric Magnetization-Prepared Rapid Gradient Echo (MPRAGE) and a T2-weighted 2D Fluid-Attenuated Inversion Recovery (FLAIR). The details of the vendor, scanner model, and scan parameters are shown in Supplementary File 1.

The neurocognitive assessment was assessed through a categorical confirmatory factor analysis^17^, with multiple tests quantifying cognitive performance across memory, language, and executive domains. Tests for the memory domain include the Delayed Word Recall Test (DWRT), Logical Memory Test (LMT), and an incidental learning test (IL); tests for the language domain include Animal Naming Test (ANT), Boston Naming Test (BNT), and a word fluency test (WFT); and tests for the executive domain include Trail Making Test Parts A and B (TMT-A, TMT-B), the Digit-Symbol Substitution Test (DSST), and the Digit Span Backward Test (DSB). The global cognitive function was calculated from all tests from the three domains, as well as item-level data from the Mini-Mental State Examination (MMSE). A lower cognitive factor score indicates poorer cognitive function within the respective domain.

1,972 participants with usable MRI scan were included in our study. Among these participants, we further excluded participants with poor quality FLAIR images (N=2), missing factor score for neurocognitive assessment (N=16), had diagnosed dementia at Visit 5 (baseline; N=110), missing genomic or vascular risk factors (N=136), and race/center categories with an insufficient number (total N=41: Black/Forsyth County, N=28; Black/ Minneapolis, N=1; Black/Washington County, N=8; other race group, N=4). The final cohort includes 1,667 participants. A flow diagram illustrating the derivation of the final analytic sample, including the number of participants excluded at each stage, is shown in Supplementary Figure 1. These participants are weighted by the ARIC-NCS sampling strategy back to Visit 5 to ensure that the sample reflects the characteristics of the entire Visit 5 cohort^15, 18^. These sampling weights were incorporated into all regression and linear mixed-effects model analyses to support inference representative of the Visit 5 cohort. Visit 5 served as the baseline reference for this study. Because loss to follow-up may not occur completely at random, we compared baseline demographic, clinical, and imaging characteristics between participants with and without longitudinal follow-up. We used chi-square tests for categorical variables and independent-samples t-tests for continuous variables, with standardized mean differences (SMD) additionally calculated for all variables to quantify the magnitude of between-group differences independent of sample size^19^.

Demographic and genomic characteristics (age, sex, race, education level, APOE carrier status) were obtained from Visit 1, and clinical characteristics of participants were measured at Visit 5. This included body mass index (BMI), hypertension, type 2 diabetes, and low-density lipoprotein (LDL) cholesterol level. Education level was categorized as less than high school, high school or GED equivalent, or at least some college. APOE carrier status was assigned as carrying none (reference) or at least one ε4 allele. Hypertension was defined as a systolic blood pressure of 140 mmHg or higher, a diastolic blood pressure of 90 mmHg or higher, or the use of antihypertensive medication, consistent with the criteria established by the Seventh Report of the Joint National Committee on Prevention, Detection, Evaluation, and Treatment of High Blood Pressure^20^. Type 2 diabetes was defined as a fasting glucose level of 126 mg/dL or higher, a non-fasting glucose level of greater than 200 mg/dL, use of diabetic medication, or self-reported diagnosis of diabetes by physician, consistent with the diabetes ascertainment criteria used in the Atherosclerosis Risk in Communities (ARIC) Study and with prior reports from this cohort^15, 21^. LDL cholesterol was recorded in SI Unit (mmol/L) and was modeled as a continuous variable in all regression analyses; no predefined categorical threshold was applied to dichotomize LDL level. Cigarette smoking status was self-reported as ever smoker or never smoker. These characteristics were included as covariates in the regression model.

### MRI measures and WMH quantification

MPRAGE images were parcellated with OpenMAP-T1 (https://github.com/OishiLab/OpenMAP-T1)^14^ and co-registered to FLAIR, on which WMH were segmented using OpenMAP_FL (https://github.com/OishiLab/OpenMAP-FL). Of 280 JHU_MNI anatomical labels^22^, 112 white matter regions were analyzed (Supplementary File 2). Regional WMH burden was calculated as WMH volume divided by the participant-specific native-space volume of each region. Total WMH burden was calculated as total WMH volume divided by participant-specific total white matter volume. Because both measures used subject-specific anatomical denominators, no additional TCV/ICV normalization was applied.

### Statistical analysis

We characterized the spatial patterns of WMH distribution using principal component analysis (PCA) as an unsupervised spatial phenotyping method. The percentage WMH volume in each of 112 anatomically defined white matter regions was converted to Z-scores prior to PCA. Regions with constant (zero) variance were dropped before PCA as they provide no discriminative information. Removing them does not affect the PCA result for the remaining regions but prevents degenerate eigenvalues that can interfere with component-retention methods such as parallel analysis. PCA was performed using baseline imaging data from Visit 5. Analyses were conducted using the prcomp function in R.

The component selection process followed a two-stage procedure. First, the number of retained candidate PCs was determined using Horn’s parallel analysis, implemented in the psych package (version 2.4.3), which identified PCs with eigenvalues exceeding those derived from randomly generated data. Second, the stability of each candidate component was evaluated using Tucker’s congruence coefficients across resampled split samples. Specifically, to assess PC stability, 80% of participant scans were randomly selected without replacement across 100 iterations, PCA was repeated in each resampled subset, and Tucker’s congruence coefficients were calculated between the subset-derived components and the corresponding components from the full dataset. Component congruence quantifies the similarity between two loading vectors and is commonly used to evaluate whether PCA-derived spatial patterns are reproducible across samples. PCs with a median Tucker’s congruence coefficient ≥0.85 across resampling iterations were considered stable and retained as the final component set for downstream analyses^23^.

To further evaluate the robustness of the retained PCA solution, two additional decompositions were compared with the original solution using Tucker’s congruence coefficient. First, a residualized PCA was performed after regressing each regional WMH percentage on total WMH percentage to assess whether retained components reflected spatial distribution rather than overall lesion burden. Second, an out-of-fold, site-wise PCA was performed to evaluate cross-site robustness: for each ARIC field center, PCA was fit using the remaining three centers, and the resulting loadings were used to project participants from the held-out center. This produced full-cohort PC scores in which no participant was scored using a solution derived from their own center. Because PCA component order and sign are arbitrary, site-specific components were matched and sign-aligned to the original solution using Tucker’s congruence coefficient. Congruence coefficients were used to assess reproducibility across sites and after removal of global WMH burden.

The associations between selected PC and demographic, genomic, and clinical covariates were evaluated using generalized linear models. Age and sex were analyzed without adjustment; analyses of other variables were adjusted for age and sex. False discovery rate (FDR) correction was applied to control for multiple comparisons.

To investigate the relationships between WMH characteristics and cognitive performance at baseline (Visit 5) and follow-up (Visits 6 and/or 7), we used linear mixed-effects models (LMM) implemented with lmerTest package in R. This approach accounts for both the cross-sectional state and the longitudinal trajectory of cognitive function while handling within-subject correlation across multiple timepoints. The model included cognitive scores at any available time point as the outcome, with participant ID as a random intercept. The fixed effects include the time interval (years from baseline), age, sex, race, education, APOE carrier status, body mass index, hypertension, diabetes, LDL cholesterol and cigarette smoking status. In the first analysis, we focused on the effect of overall WMH burden on cognitive performance by including total WMH volume and its interaction with the time interval as independent variables. In the second analysis, we investigated the impact of WMH distribution patterns on cognitive performance and subsequent cognitive decline. Accordingly, total WMH volume was replaced with each PC score and the interaction term between the PC score and the time interval. The interaction term evaluates whether baseline WMH burden or spatial pattern is associated with the subsequent rate of cognitive change. An overview of study methodology is shown in Figure 1. As a sensitivity analysis, we repeated the linear mixed-effects models for the PC measures with additional adjustment for total WMH percentage to assess whether associations between WMH spatial patterns and cognitive decline were independent of overall WMH burden.

**Figure 1.**
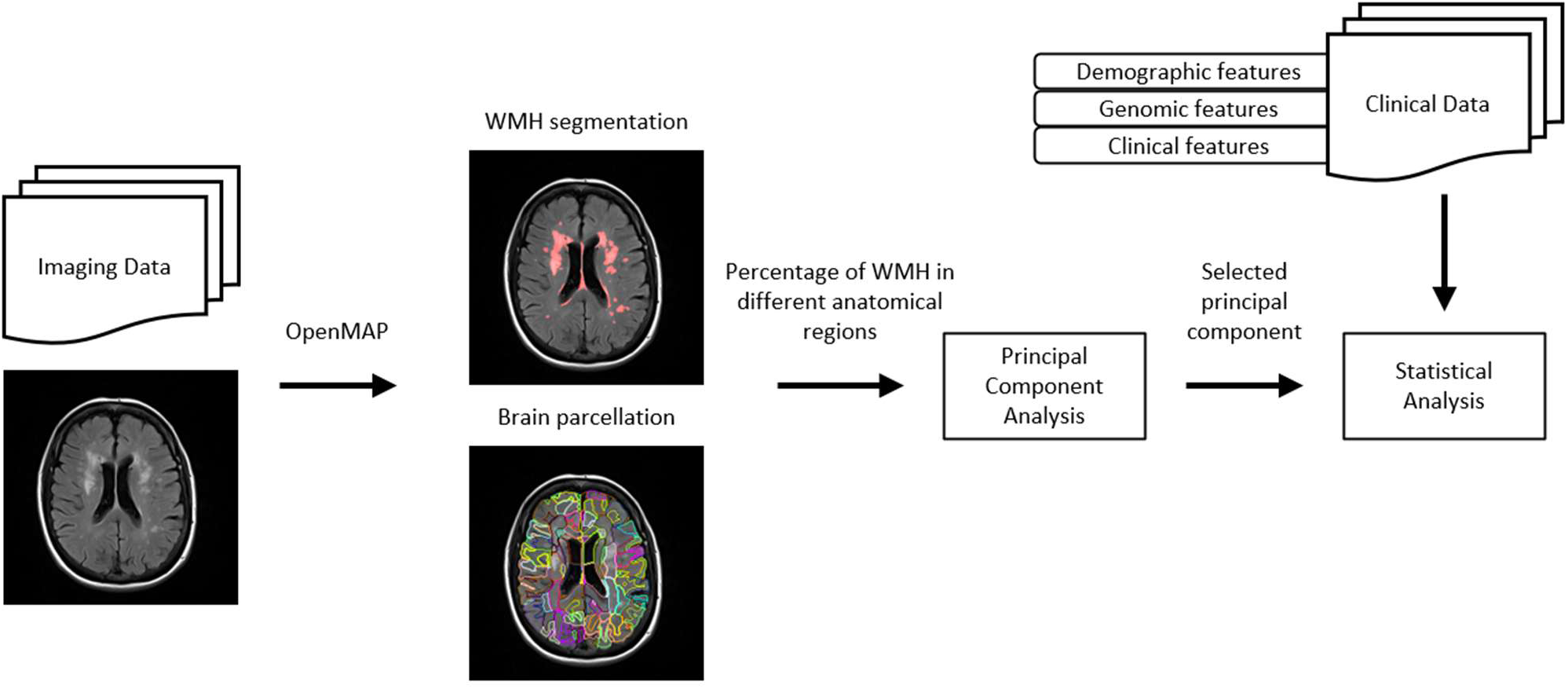
Study Methodology.

To assess robustness to selective loss to follow-up, we repeated the longitudinal LMMs using inverse probability of attrition weights. The probability of having longitudinal follow-up was estimated using logistic regression based on measured baseline characteristics associated with follow-up status, and the inverse of this estimated probability was used to weight observed participants in the sensitivity analysis. To evaluate the spatial and cross-site robustness of the PC-derived associations, we further repeated the LMMs using PC scores from the residualized PCA and from the out-of-fold, site-wise PCA described above.

Imaging analyses were performed in Python (version 3.11.14), and statistical analyses were performed in R (version 4.3.3).

## Results

The demographic and clinical characteristics of the participants at baseline (Visit 5) are shown in Table 1. Among the 1,667 participants, 1,135 (68.09%) had follow up cognitive assessment at Visit 6 (time interval Mean = 4.88, SD = 0.61 years), 1,052 (63.11%) had follow up cognitive assessment at Visit 7 (time interval Mean = 6.54, SD = 0.66 years), 957 (57.41%) had cognitive assessment at both visits.

**Table 1.** Participant demographic and clinical characteristics.

|  | Overall<br>(N=1667) | One Follow Up<br>Visit (N=273) | Two Follow Up<br>Visit (N=957) |
| --- | --- | --- | --- |
| Baseline Age (years) | 76.2 (5.26) | 76.9 (5.22) | 74.9 (4.76) |
| Sex |  |  |  |
| Female | 993 (59.6%) | 156 (57.1%) | 565 (59.0%) |
| Male | 674 (40.4%) | 117 (42.9%) | 392 (41.0%) |
| Education Level |  |  |  |
| Less than high school | 217 (13.0%) | 37 (13.6%) | 102 (10.7%) |
| High school graduate or GED equivalent | 703 (42.2%) | 123 (45.1%) | 371 (38.8%) |
| At least some college | 747 (44.8%) | 113 (41.4%) | 484 (50.6%) |
| Race - Study Center |  |  |  |
| White - F | 393 (23.6%) | 55 (20.1%) | 229 (23.9%) |
| Black - J | 450 (27.0%) | 74 (27.1%) | 267 (27.9%) |
| White - M | 385 (23.1%) | 71 (26.0%) | 218 (22.8%) |
| White - W | 439 (26.3%) | 73 (26.7%) | 243 (25.4%) |
| APOE ε4 Carrier | 472 (28.3%) | 91 (33.3%) | 262 (27.4%) |
| BMI (kg/m<sup>2</sup>) | 28.4 (5.63) | 28.8 (5.84) | 28.8 (5.54) |
| LDL Cholesterol (mmol/L) | 2.72 (0.887) | 2.75 (0.910) | 2.74 (0.855) |
| Diabetes | 541 (32.5%) | 101 (37.0%) | 285 (29.8%) |
| Hypertension | 1245<br>(74.7%) | 212 (77.7%) | 688 (71.9%) |
| Ever Smoker | 901 (54.0%) | 157 (57.5%) | 514 (53.7%) |
| Total WMH percentage | 5.86 (6.24) | 6.11 (6.40) | 5.20 (5.50) |

Data shown in mean (standard deviation) for continuous variables, and count (percentage) for categorical variables. Total WMH percentage represents WMH volume relative to total white matter volume. Study centers: F, Forsyth County, North Carolina; J, Jackson, Mississippi; M, Minneapolis, Minnesota; W, Washington County, Maryland. Abbreviations: APOE ε4, Apolipoprotein E epsilon 4; BMI, body mass index; LDL, low-density lipoprotein; WMH, white matter hyperintensities.

A total of 73% of participants had at least one follow-up evaluation, with a mean (SD) baseline-to-follow-up interval of 6.3 (0.86) years. Baseline characteristics of participants with and without longitudinal follow-up are compared in Supplementary Table 1. Participants without follow-up were older, had lower educational attainment, higher BMI, higher LDL cholesterol, a higher prevalence of hypertension, and greater total WMH burden at baseline (all p<0.05, |SMD|≥0.1).

Using the OpenMAP deep learning-based structural parcellation and WMH segmentation algorithm, we generated % WMH volumes for 112 white matter regions. Thirteen regions with zero variance across all participants were excluded from analysis, leaving 99 regions for principal component analysis. Horn’s parallel analysis identified 14 candidate principal components (PCs) that explained more variance than would be expected by chance, based on the scree plot shown in Supplementary Figure 2. Of these 14 candidates, Tucker’s congruence analysis across resampled split samples supported the stability of the first eight PCs, with median congruence coefficients ≥0.85 across 100 resampling iterations. Congruence coefficients for PC9 and subsequent components were <0.85, indicating reduced stability across subsamples (Supplementary Figure 3). The eight stable components were therefore retained as the final component set for downstream analyses. Visualizations of the spatial loading patterns for the retained PCs are shown in Figure 2. Details of the loading and explained variance are shown in Supplementary File 3 and Supplementary File 4.

**Figure 2.**
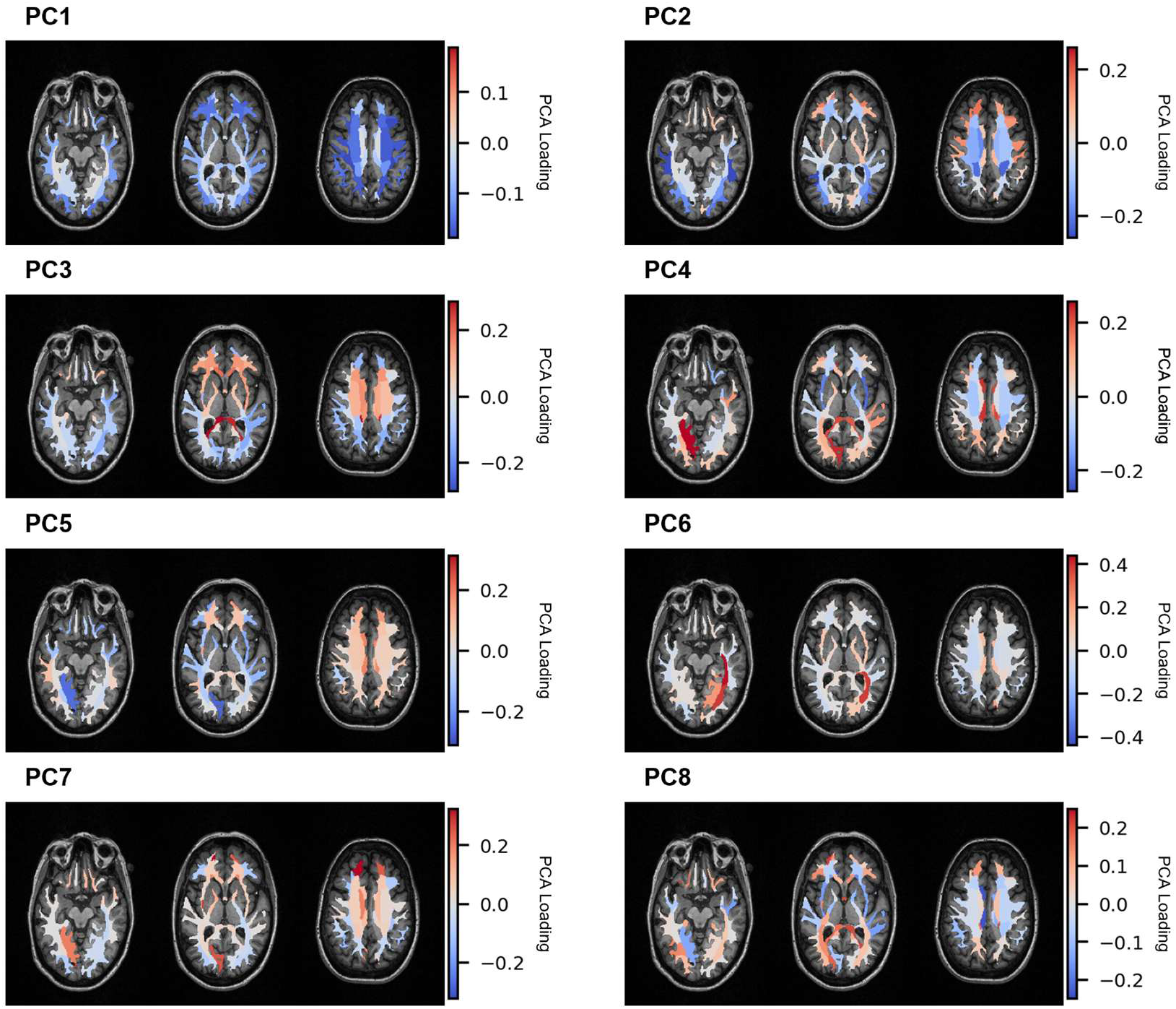
Principal component loading by parcellation. *Loadings for selected principal components are overlaid on representative axial brain slices for visualization. Colors denote the sign and magnitude of regional loadings (red = positive; blue = negative; white/gray ≈ 0), with color bars indicating loading values for each PC. Regions with larger absolute loadings contribute more strongly to the corresponding WMH spatial pattern*.

The first PC explained 20.46% of total variance, with widespread, bilateral negative loadings across regions (Table 2). The highest contributing regions include the superior longitudinal fasciculus and the superficial white matter areas just beneath the middle frontal gyrus, the postcentral and precentral gyri, and the angular gyrus. From the second PC onward, explained variance dropped substantially to 4.50%, with loadings reflecting more localized WMH patterns. Together, the eight PCs explained 42.10% of cumulative variance.

**Table 2.** Spatial Distribution of White Matter Hyperintensity of Selected Principal Component.

| Principal Components | Characteristic of White Matter Hyperintensity Distribution | Total Variance Explained (%) |
| --- | --- | --- |
| PC1 | Global negative burden pattern, particularly in the association fibers | 20.46 |
| PC2 | Frontal-dominant superficial white matter burden with preserved temporoparietal areas | 4.59 |
| PC3 | Deep white matter burden with preserved superficial white matter | 3.59 |
| PC4 | Posterior white matter and cingulum bundle burden | 3.20 |
| PC5 | Negative burden in the internal capsule, middle frontal, temporal, and medial parietal areas with preserved superior white matter | 3.08 |
| PC6 | Burden localized to the left posterior thalamic radiation | 2.50 |
| PC7 | Superior frontal, periventricular, and posterior medial white matter burden | 2.40 |
| PC8 | Burden in limbic and limbic-related fibers | 2.28 |

Distinct spatial patterns of WMH were differentially associated with demographic, clinical, and vascular risk factors (Table 3 and Supplementary Figure 4). PC1 showed a strong inverse correlation with total WMH volume, suggesting that it represents a global WMH burden factor, with lower scores indicating higher overall WMH load. Older age, higher BMI, and hypertension were significantly associated with increased overall WMH burden. PC2, characterized by positive loadings on anterior frontal and pericentral white matter and negative loadings on posterior temporal-occipital-parietal white matter, was significantly positively correlated with age. PC3, defined by high positive loadings in the splenium and genu of the corpus callosum as well as the anterior corona radiata, exhibited significant positive associations with female sex and diabetes. PC4, with positive loadings on the splenium of the corpus callosum and cingulate gyrus and negative loadings on the external capsule and frontal white matter as main contributors, was significantly associated with the Black race–Jackson study-center category. Additional sex-specific patterns emerged in PC7. No significant associations were observed with LDL cholesterol, education, APOE ε4 carrier status, or smoking history after FDR correction.

**Table 3.** Correlation between Principal Component and Demographic, Clinical and Genomic Factors.

| PC | Age | Total WMH | Sex (Male) | Race (White) | BMI | LDL | Education | APOE4 | Diabetes | Hypertension | Ever Smoker |
| --- | --- | --- | --- | --- | --- | --- | --- | --- | --- | --- | --- |
| PC1 | -0.2255<br>(<0.0001) | -0.9697<br>(<0.0001) | 0.0579<br>(0.1269) | 0.0601<br>(0.1153) | -0.0741<br>(0.0293) | 0.0154<br>(0.8082) | 0.0210<br>(0.7270) | 0.0243<br>(0.6799) | -0.0122<br>(0.8502) | -0.1032<br>(0.0005) | -0.0025<br>(0.9742) |
| PC2 | 0.1066<br>(0.0003) | 0.0411<br>(0.3889) | 0.0453<br>(0.3418) | 0.0344<br>(0.5158) | 0.0178<br>(0.7666) | -0.0353<br>(0.5030) | 0.0283<br>(0.6126) | -0.0292<br>(0.5988) | 0.0414<br>(0.3889) | 0.0149<br>(0.8092) | -0.0091<br>(0.8601) |
| PC3 | 0.0650<br>(0.0683) | -0.1152<br>(0.0001) | -0.2319<br>(<0.0001) | 0.0106<br>(0.8502) | -0.0012<br>(0.9766) | -0.0360<br>(0.4977) | 0.0166<br>(0.7910) | -0.0021<br>(0.9742) | 0.0892<br>(0.0041) | 0.0213<br>(0.7270) | 0.0148<br>(0.8092) |
| PC4 | 0.1106<br>(0.0002) | 0.0237<br>(0.6963) | 0.0069<br>(0.8942) | -0.0883<br>(0.0043) | -0.0110<br>(0.8502) | 0.0374<br>(0.4779) | -0.0329<br>(0.5335) | 0.0475<br>(0.3244) | 0.0341<br>(0.5158) | 0.0150<br>(0.8092) | 0.0171<br>(0.7783) |
| PC5 | -0.0228<br>(0.7001) | 0.1042<br>(0.0004) | -0.0590<br>(0.1174) | -0.0008<br>(0.9795) | 0.0416<br>(0.3889) | -0.0022<br>(0.9742) | -0.0337<br>(0.5200) | -0.0230<br>(0.7001) | 0.0399<br>(0.4076) | 0.0111<br>(0.8502) | -0.0459<br>(0.3358) |
| PC6 | -0.0171<br>(0.7783) | -0.0431<br>(0.3791) | -0.0103<br>(0.8502) | 0.0140<br>(0.8344) | -0.0204<br>(0.7361) | 0.0019<br>(0.9742) | 0.0099<br>(0.8548) | 0.0125<br>(0.8502) | -0.0202<br>(0.7361) | -0.0032<br>(0.9742) | -0.0295<br>(0.5988) |
| PC7 | -0.0666<br>(0.0670) | -0.0391<br>(0.4276) | -0.0827<br>(0.0092) | 0.0284<br>(0.6126) | -0.0087<br>(0.8610) | -0.0021<br>(0.9742) | 0.0304<br>(0.5916) | -0.0107<br>(0.8502) | -0.0006<br>(0.9795) | -0.0117<br>(0.8502) | -0.0107<br>(0.8502) |
| PC8 | 0.0423<br>(0.3889) | -0.0366<br>(0.4838) | 0.0199<br>(0.7361) | -0.0313<br>(0.5648) | 0.0417<br>(0.3889) | -0.0114<br>(0.8502) | 0.0502<br>(0.2602) | 0.0019<br>(0.9742) | 0.0181<br>(0.7641) | -0.0200<br>(0.7361) | 0.0160<br>(0.8016) |

*Correlations with age and sex were unadjusted. All other correlations were adjusted for age and sex using partial correlation. Data shown in coefficient (p-value), all p-values were corrected for multiple testing using the False Discovery Rate.*

Associations between baseline WMH patterns and cognitive performances at Visit 5, derived from the main effects of the linear mixed-effects model, are shown in Table 4. Total WMH percentage, as well as PC1, was significantly associated with cognitive performance in global cognition, language, and executive function, but not in memory. Among the spatially specific patterns, PC8, characterized by high positive loadings on the fornix, splenium of the corpus callosum, and occipital white matter, was significantly associated with poorer baseline executive function.

**Table 4.** Linear Mixed-Effects Model Estimates for Main Effects of WMH Patterns on Cognitive Performance.

|  | Global | Executive | Language | Memory |
| --- | --- | --- | --- | --- |
| Total WM % | -0.0106[-0.0159, -0.0055]<br>($<0.0001$) | -0.0114[-0.0172, -0.0056]<br>(0.0001) | -0.0098[-0.0158, -0.0039]<br>(0.0011) | -0.0040[0.0102, 0.0021]<br>(0.1992) |
| PC1 | 0.0150[0.0077, 0.0222]<br>(0.0003) | 0.0165[0.0084, 0.0246]<br>(0.0010) | 0.0136[0.0054, 0.0219]<br>(0.0096) | 0.0053[-0.0033, 0.0138]<br>(0.6010) |
| PC2 | -0.0062[-0.0214, 0.0090]<br>(0.4558) | -0.0101[-0.0272, 0.0069]<br>(0.3548) | -0.0111[-0.0285, 0.0062]<br>(0.4771) | -0.0059[-0.0237, 0.0119]<br>(0.6887) |
| PC3 | 0.0084[-0.0092, 0.0259]<br>(0.4403) | 0.0099[-0.0096, 0.0294]<br>(0.3946) | 0.0052[-0.0148, 0.0251]<br>(0.6848) | 0.0112[-0.0093, 0.0318]<br>(0.6173) |
| PC4 | 0.0084[-0.0095, 0.0263]<br>(0.4403) | 0.0027[-0.0171, 0.0225]<br>(0.7865) | 0.0057[-0.0145, 0.0260]<br>(0.6848) | 0.0180[-0.0029, 0.0389]<br>(0.2920) |
| PC5 | 0.0173[-0.0007, 0.0353]<br>(0.1901) | 0.0218[0.0020, 0.0415]<br>(0.0988) | 0.0113[-0.0090, 0.0315]<br>(0.5495) | 0.0077[-0.0133, 0.0287]<br>(0.6842) |
| PC6 | -0.0108[-0.0325, 0.0109]<br>(0.4403) | -0.0128[-0.0374, 0.0117]<br>(0.3946) | -0.0235[-0.0492, 0.0022]<br>(0.2325) | 0.0018[-0.0241, 0.0277]<br>(0.8952) |
| PC7 | 0.0130[-0.0079, 0.0339]<br>(0.3568) | 0.0072[-0.0160, 0.0304]<br>(0.5782) | 0.0119[-0.0120, 0.0358]<br>(0.5569) | -0.0091[-0.0337, 0.0155]<br>(0.6842) |
| PC8 | -0.0180[-0.0394, 0.0035]<br>(0.2312) | -0.0300[-0.0538, -0.0061]<br>(0.0416) | -0.0167[-0.0413, 0.0078]<br>(0.4771) | 0.0017[-0.0235, 0.0269]<br>(0.8952) |

*Data shown in coefficient [95% Confidence Interval] (p-value), all p-values for PCs were corrected for multiple testing using the False Discovery Rate within each cognitive domain. The main effects reflect the cross-sectional association at baseline (time interval = 0). Total WM %: total white matter percentage relative to white matter volume. PC: principal component.*

The longitudinal impacts of WMH patterns on the rate of cognitive decline, assessed via the interaction terms, are presented in Table 5. Greater total WMH burden was significantly associated with accelerated cognitive decline across all four domains: global cognition, executive function, language, and memory. PC3, characterized by greater deep white matter burden involving the corpus callosum and anterior corona radiata with relative preservation of superficial white matter, was significantly associated with accelerated decline in global cognition, language, and memory despite showing no significant cross-sectional association at baseline.

**Table 5.** Linear Mixed-Effects Model Estimates for Longitudinal Interaction Effects on Cognitive Performance.

|  | Global | Executive | Language | Memory |
| --- | --- | --- | --- | --- |
| Total WM %:Time | -0.0016[-0.0023, -0.0010]<br>($<0.0001$) | -0.0017[-0.0025, -0.0008]<br>(0.0001) | -0.0012[-0.0022, -0.0003]<br>(0.0082) | -0.0014[-0.0022, -0.0006]<br>(0.0011) |
| PC1:Time | 0.0022[0.0014, 0.0031]<br>($<0.0001$) | 0.0021[0.0009, 0.0033]<br>(0.0031) | 0.0020[0.0007, 0.0032]<br>(0.0100) | 0.0018[0.0007, 0.0030]<br>(0.0302) |
| PC2:Time | -0.0016[-0.0035, 0.0002]<br>(0.2140) | -0.0019[-0.0044, 0.0005]<br>(0.2797) | 0.0027[0.0001, 0.0053]<br>(0.1673) | -0.0023[-0.0047, 0.0002]<br>(0.2794) |
| PC3:Time | -0.0043[-0.0063, -0.0022]<br>(0.0003) | -0.0019[-0.0047, 0.0009]<br>(0.3507) | -0.0051[-0.0080, -0.0022]<br>(0.0096) | -0.0039[-0.0066, -0.0011]<br>(0.0457) |
| PC4:Time | -0.0017[-0.0039, 0.0005]<br>(0.2489) | 0.0013[-0.0017, 0.0042]<br>(0.4571) | 0.0007[-0.0024, 0.0038]<br>(0.6848) | -0.0030[-0.0059, -0.0001]<br>(0.2366) |
| PC5:Time | -0.0014[-0.0037, 0.0008]<br>(0.3568) | -0.0037[-0.0066, -0.0007]<br>(0.0616) | 0.0011[-0.0021, 0.0043]<br>(0.6651) | 0.0003[-0.0026, 0.0033]<br>(0.8952) |
| PC6:Time | -0.0001[-0.0028, 0.0027]<br>(0.9553) | 0.0022[-0.0015, 0.0058]<br>(0.3548) | 0.0004[-0.0035, 0.0043]<br>(0.8378) | -0.0017[-0.0053, 0.0020]<br>(0.6514) |
| PC7:Time | 0.0011[-0.0016, 0.0037]<br>(0.4558) | 0.0029[-0.0005, 0.0064]<br>(0.2575) | 0.0014[-0.0023, 0.0051]<br>(0.6651) | 0.0018[-0.0017, 0.0053]<br>(0.6173) |
| PC8:Time | -0.0028[-0.0053, -0.0003]<br>(0.1169) | -0.0020[-0.0054, 0.0014]<br>(0.3548) | -0.0017[-0.0053, 0.0019]<br>(0.5569) | -0.0007[-0.0040, 0.0027]<br>(0.8477) |

Data shown in coefficient [95% Confidence Interval] (p-value), all p-values for PCs were corrected for multiple testing using the False Discovery Rate within each cognitive domain. Total WM %: total white matter percentage relative to white matter volume. PC: principal component.

To evaluate whether the two spatially specific components identified above (PC3 and PC8) were robust, we compared the original PCA solution against a residualized PCA and an out-of-fold, site-wise PCA using Tucker’s congruence coefficient (Supplementary Figure 5). In the residualized PCA, PC3 showed high congruence with the original solution (best-match φ = 0.95), while PC8 showed lower congruence (best-match φ = 0.61). In the out-of-fold, site-wise PCA, in which each participant was scored using a solution derived exclusively from the other three field centers, both PC3 and PC8 showed consistently high congruence with the original solution across all four held-out centers (PC3: mean φ = 0.89, range 0.77–0.98; PC8: mean φ = 0.91, range 0.87–0.96).

Since participants without longitudinal follow-up differed from those with follow-up on several baseline characteristics (Supplementary Table 1), we repeated the linear mixed-effects models using inverse probability of attrition weights (Supplementary Table 2). The PC3-by-time interaction remained significantly associated with accelerated decline in global cognition, language, and memory after attrition weighting. The PC1-by-time interaction remained significant across all four domains.

In sensitivity analyses additionally adjusting for total WMH percentage, the associations involving spatially specific components were largely unchanged (Supplementary Table 3). The PC3-by-time interaction remained significantly associated with accelerated decline in global cognition and language, with a comparable estimate in memory that approached significance. The cross-sectional association between PC8 and baseline executive function also persisted after adjustment. The cross-sectional association between PC1 and baseline cognitive performance was attenuated after adjustment for total WMH percentage, consistent with the strong correlation between PC1 and overall WMH burden (r = −0.97); the longitudinal PC1-by-time interaction, in contrast, remained significant across all four domains.

We further repeated the models using PC scores derived from the two alternative decompositions method. In the residualized PCA (Supplementary Table 4), in which the linear effect of total WMH volume was removed from each region prior to decomposition, the PC3-by-time interaction likewise remained significantly associated with decline in global cognition and language, with the memory estimate remaining in the same direction. In the out-of-fold, site-wise PCA (Supplementary Table 5), the PC3-by-time interaction showed estimates consistent in direction and magnitude with the primary analysis across global cognition, language, and memory, and the PC1-by-time interaction remained significant across all four domains. Neither alternative decomposition reproduced the cross-sectional association between PC8 and baseline executive function.

## Discussion

In this study, we investigated the relationship between WMH distribution patterns and cognitive function in an older population using a combination of structural parcellation and WMH lesion segmentation. WMH was quantified continuously across 112 anatomically defined white matter regions in a large community-based cohort with approximately six years of cognitive follow-up, allowing spatial distribution to be characterized without recourse to a priori territorial divisions or post hoc lesion clusters. This approach provided anatomically precise localization of pathological changes. Principal component analysis identified 14 candidate components, of which eight met our resampling-based stability criteria and were retained as reproducible spatial patterns of WMH distribution.

Correlation analysis revealed that these patterns were differentially associated with established WMH risk factors and demographic characteristics (Table 3). PC1 showed a strong inverse correlation with total WMH volume, confirming its role as a marker for overall WMH severity. The significant associations with older age, higher BMI, and hypertension align with established vascular risk factors for WMH accumulation^24–26^. The association between PC3 (deep WMH burden) and diabetes was also observed in other cohort studies^27, 28^. One possible explanation is that diabetes-related small-vessel injury preferentially affects deep perforating vascular territories^29^; however, this mechanism was not directly evaluated in the present study. The remaining components showed associations with age, sex, and race–study center (Table 3) but were not accompanied by cognitive associations. Race and recruitment site are perfectly correlated and completely confounded in these analyses, limiting our ability to independently attribute these effects to race versus regional environmental factors. Future studies with more geographically diverse cohorts are needed to disentangle these variables. Notably, we found no significant association between any WMH pattern and APOE ε4 carrier status.

Several WMH measures were associated with cognitive outcomes, although the observed associations were not uniformly domain specific. Global WMH burden was significantly associated with poorer performance in global cognition, language, and executive function at baseline, and was also a significant marker of accelerated cognitive decline across all four domains (global, executive, language, and memory) during the follow-up period. PC1 correlated very strongly with total WMH burden (r = −0.97), indicating that it largely represents global lesion severity rather than an independent spatial distribution pattern. Consistent with this interpretation, its cross-sectional cognitive associations were attenuated after adjustment for total WMH burden.

PC3, characterized by deep white matter burden within the corpus callosum and anterior corona radiata, with relatively preserved superficial white matter, showed no significant cross-sectional associations at baseline but emerged as a significant marker of accelerated cognitive decline in global cognition, language, and memory. The anterior corona radiata and corpus callosum serve as critical pathways for thalamocortical^30, 31^ and interhemispheric communication^32, 33^, respectively. As these pathways support widely distributed cortical networks, their disruption would affect multiple cognitive domains. One possible explanation for the absence of a baseline association is that relatively preserved superficial pathways may partially compensate for deep white matter injury ^34, 35^. This interpretation is speculative, however, and cannot be established from the present cross-sectional imaging data.

PC8 showed a cross-sectional association with executive function but was not associated with the rate of subsequent cognitive change. Although the spatial PC8 pattern was reproducible across the site-wise decompositions, it showed low congruence with the residualized PCA solution, and its cross-sectional executive-function association was not reproduced in the residualized or out-of-fold models, or in the sensitivity analysis with inverse probability attrition weighting. We therefore regard this finding as exploratory and requiring independent replication.

To assess whether the longitudinal associations were affected by selective attrition, the mixed-effects models were reweighted by the inverse probability of having follow-up data. These WMH patterns and their time interaction remained significant, with estimates comparable to the primary analysis (Supplementary Table 2).

Sensitivity analyses further supported the robustness of the PC3 findings. Adjustment for total WMH percentage, and repeating the decomposition on regional values residualized for total WMH volume, left the PC3-by-time interaction essentially unchanged, indicating that the association is not a function of overall lesion burden. In the out-of-fold analysis, where each ARIC field center was scored using a solution derived exclusively from the other three centers, PC3 was recovered across all four folds (mean |φ| = 0.89; range 0.77–0.98) and its association with subsequent decline was preserved. Notably, congruence with the original solution was lowest when the Jackson center was held out (φ = 0.77, versus 0.86–0.98 for the remaining folds); because all Black participants in this cohort were recruited at Jackson and the remaining centers enrolled White participants, this fold differs from the others in both site and race, and the reduced congruence cannot be attributed to either factor alone. Despite this variability, effect estimates for the PC3-by-time interaction were consistent in direction and magnitude with the primary models across sensitivity analyses (Supplementary Tables 2–5).

This study has several limitations. WMH segmentation from FLAIR images may be susceptible to various technical and biological confounds. Motion artifacts, enlarged perivascular spaces, lacunes, cystic lesions, infarcts, and other non-WMH hyperintense or hypointense abnormalities can influence automated segmentation accuracy. Although images underwent quality control procedures and cases with poor-quality FLAIR scans were excluded from analysis, subtle artifacts or lesion mimics may have remained undetected and influenced regional WMH estimates, particularly in smaller anatomical regions where such errors would constitute a larger proportional impact. Such measurement errors would be expected to introduce noise into the PCA-derived spatial patterns and could potentially attenuate the observed associations with cognitive outcomes, suggesting that the true effect sizes may be underestimated. Principal component analysis decomposes covariance among regional WMH measurements and yields patterns that are defined statistically rather than biologically. The components identified here were not validated against independent mechanistic, physiological, or neuropathological measures, and whether they correspond to distinct underlying disease processes remains unknown. Our cohort consisted only of Black and White participants, which limited its generalizability to other racial and ethnic groups. Moreover, because race and recruitment site are perfectly correlated and completely confounded, we could not separate race-associated differences from site-specific factors. Unmeasured cofounders may also have influenced the observed associations in the cohort. Selective attrition is a further limitation. Participants without follow-up were older and had greater vascular risk and total WMH burden at baseline, so the longitudinal analyses were conducted in a comparatively healthier subset. Although estimates were essentially unchanged after inverse probability of attrition weighting, weighting can only adjust for measured baseline characteristics, and residual bias from unmeasured determinants of dropout cannot be excluded. Future studies should prioritize diverse recruitment to enhance representativeness. Additionally, while all data derive from a single study with a common imaging protocol, variations in MRI scanners across study centers could introduce bias. Although the internal resampling and cross-site analysis supported the stability of the retained PCA-derived WMH patterns, external validation in independent cohorts with comparable imaging, segmentation, and cognitive assessment protocols will be necessary to confirm their generalizability. Lastly, while we demonstrated cross-sectional and longitudinal associations between WMH patterns and cognitive function, these analyses establish only population-level associations. Future work, such as confirmation in studies with longitudinal imaging, is needed to assess the predictive utility of these patterns for individual-level risk stratification.

In conclusion, continuous quantification of WMH across anatomically defined white matter regions identified a deep spatial pattern involving the corpus callosum and anterior corona radiata, with relative preservation of superficial white matter, that was associated with subsequent cognitive decline despite the absence of a cross-sectional association at baseline. This association remained consistent across analyses accounting for overall WMH burden, study center, and selective attrition, supporting further investigation of spatially resolved WMH measures in longitudinal and externally validated cohorts.

## Supporting information

Full Supplementary

## Acknowledgement

The Atherosclerosis Risk in Communities Study is carried out as a collaborative study supported by National Heart, Lung, and Blood Institute contracts (75N92022D00001, 75N92022D00002, 75N92022D00003, 75N92022D00004, 75N92022D00005). The ARIC Neurocognitive Study is supported by U01HL096812, U01HL096814, U01HL096899, U01HL096902, and U01HL096917 from the NIH (NHLBI, NINDS, NIA and NIDCD). The authors thank the staff and participants of the ARIC study for their important contributions.

R.F. Gottesman was supported by the NINDS Intramural Research Program. The contributions of the NIH author were made as part of their official duties as NIH federal employees, are in compliance with agency policy requirements, and are considered Works of the United States Government. However, the findings and conclusions presented in this paper are those of the author and do not necessarily reflect the views of the NIH or the U.S. Department of Health and Human Services.

## Author Contributions

KO and XC contributed to the conception and design of the study; XC, KN, ST, JC, YU, BAW, RFG, EG, and BGW contributed to the acquisition and analysis of data; XC and KO contributed to drafting the text or preparing the figures.

## Data Availability

The ARIC data used in this study are available subject to ARIC data use agreements and approval procedures. The image-processing tools used for structural parcellation and WMH segmentation are publicly available on GitHub, including OpenMAP-T1 (https://github.com/OishiLab/OpenMAP-T1) and OpenMAP-FL (https://github.com/ OishiLab/OpenMAP-FL).

## Use of Artificial Intelligence

No artificial intelligence or large language models were used in the design and execution of this research. AI-assisted tools were used for grammar and syntax checking of the manuscript text (ChatGPT 5.6 Terra). AI-assisted tools were also used in a limited capacity during data analysis for coding support, such as in-line autocompletion, syntax suggestions, and troubleshooting (Claude Sonnet 5). The study design, analytical logic, and algorithm implementation were developed and written by the authors; all AI-suggested code was reviewed and validated before incorporation. All scientific content, data interpretation, and conclusions are the authors’ own.

