## Supplementary material for "Spatial Patterns of White Matter Hyperintensities and Cognitive Function in Older Adults: A Community-Based Study": Full Supplementary

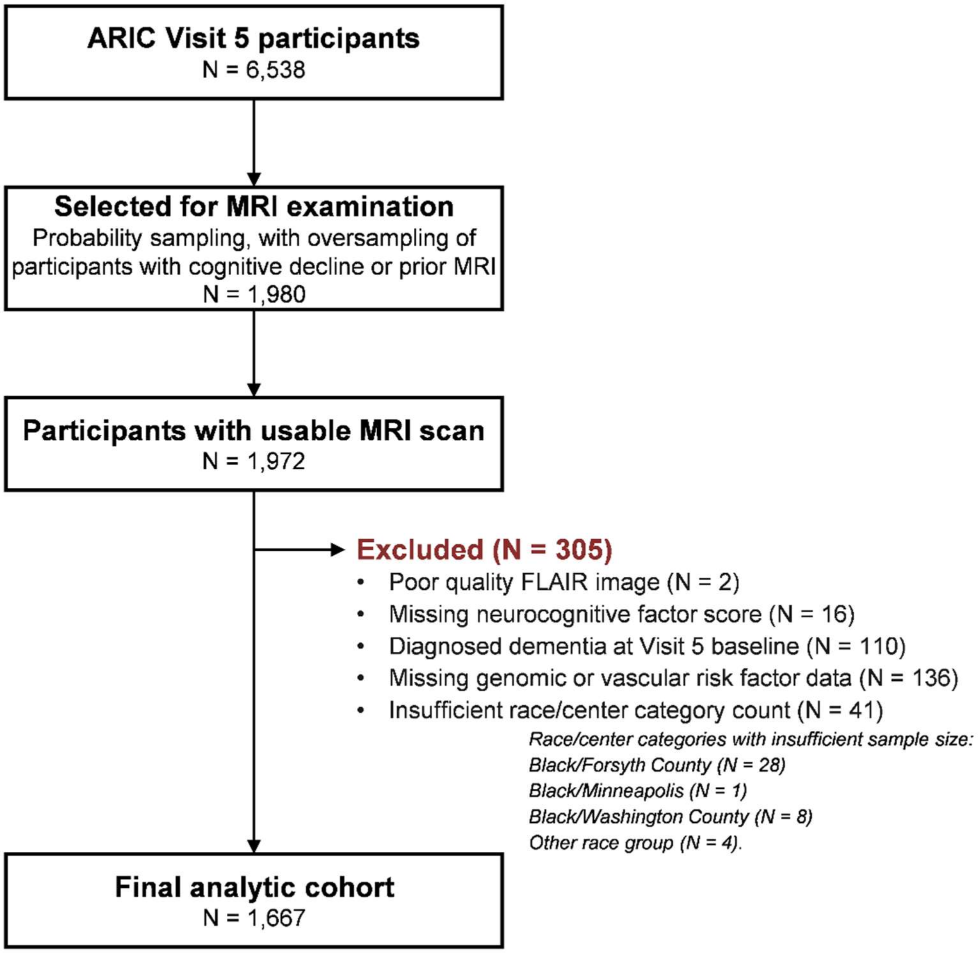


**Supplementary Figure 1**. Flow Diagram for Study Analytic Cohort

**Supplementary Table 1**: Baseline Characteristics by Longitudinal Follow-up Status

| | No Follow-up (N=437) | With Follow-up (N=1230) | p value | SMD |
|---|---|---|---|---|
| Baseline Age (years) | 78.5 (5.43) | 75.3 (4.93) | <0.001 | -0.61 |
| Sex | | | | |
| Female | 272 (62.2%) | 721 (58.6%) | 0.185 | -0.074 |
| Male | 165 (37.8%) | 509 (41.4%) | | |
| Education Level | | | | |
| Less than high school | 78 (17.8%) | 139 (11.3%) | | |
| High school graduate or GED equivalent | 209 (47.8%) | 494 (40.2%) | <0.001 | 0.306 |
| At least some college | 150 (34.3%) | 597 (48.5%) | | |
| Race - Study Center | | | | |
| White - F | 109 (24.9%) | 284 (23.1%) | | |
| Black - J | 109 (24.9%) | 341 (27.7%) | 0.483 | 0.087 |
| White - M | 96 (22.0%) | 289 (23.5%) | | |
| White - W | 123 (28.1%) | 316 (25.7%) | | |
| APOE ε4 Carrier | 119 (27.2%) | 353 (28.7%) | 0.558 | -0.033 |
| BMI (kg/m$^2$) | 27.5 (5.58) | 28.8 (5.61) | <0.001 | 0.233 |
| LDL Cholesterol (mmol/L) | 2.64 (0.938) | 2.75 (0.867) | 0.0139 | 0.111 |
| Diabetes | 155 (35.5%) | 386 (31.4%) | 0.117 | 0.087 |
| Hypertension | 345 (78.9%) | 900 (73.2%) | 0.017 | 0.136 |
| Ever Smoker | 230 (52.6%) | 671 (54.6%) | 0.489 | -0.039 |
| Total WMH percentage | 7.16 (7.35) | 5.41 (5.72) | <0.001 | -0.266 |

*Data shown in mean (standard deviation) for continuous variables, and count (percentage) for categorical variables. SMD: standardized mean difference. Total WMH percentage represents WMH volume relative to total white matter volume. Study centers: F, Forsyth County, North Carolina; J, Jackson, Mississippi; M, Minneapolis, Minnesota; W, Washington County, Maryland. Abbreviations: APOE ε4, Apolipoprotein E epsilon 4; BMI, body mass index; LDL, low-density lipoprotein; WMH, white matter hyperintensities.*

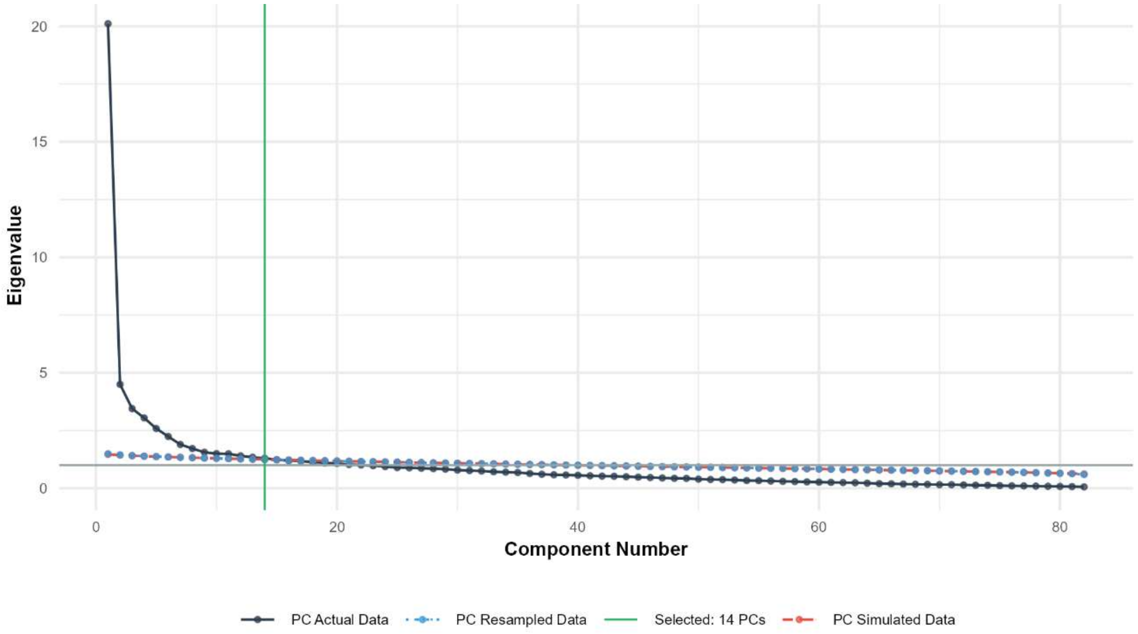


**Supplementary Figure 2**. Scree Plot of Parallel Analysis

*The scree plot displays eigenvalues across component numbers to determine the optimal number of principal components (PCs) to retain. Three data series are shown: (1) PC Actual Data (solid black line) represents eigenvalues derived from the original dataset; (2) PC Simulated Data (dashed red line) represents eigenvalues from randomly generated datasets with the same dimensions as the original data, assuming no underlying structure; and (3) PC Resampled Data (dotted blue line) represents eigenvalues from permuted versions of the original data, where values within each variable are randomly shuffled to destroy correlations while preserving distributional properties. Components are retained when actual eigenvalues exceed both simulated and resampled eigenvalues, indicating that the component captures meaningful variance beyond chance. The vertical green line indicates the number of components selected based on this criterion (n = 14 PCs). The horizontal gray line marks the Kaiser criterion threshold (eigenvalue = 1), a traditional but often liberal retention rule.*

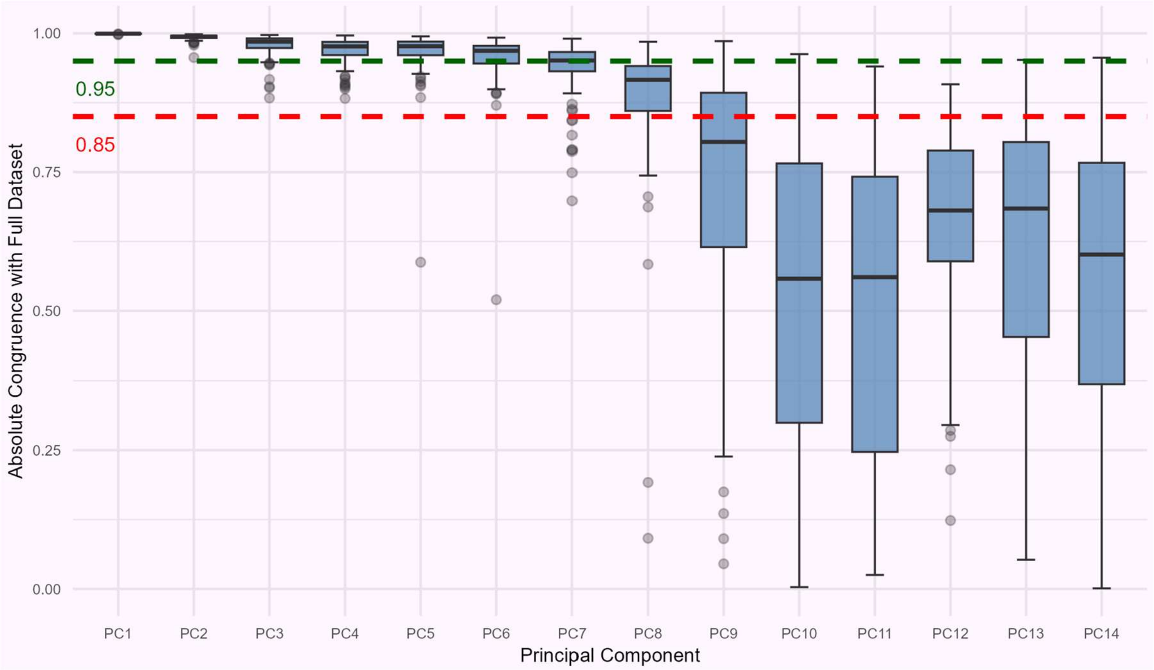


**Supplementary Figure 3**: Stability of Principal Components in Resampling Analysis

*Box plots show the absolute congruence coefficients between principal components (PCs) derived from the full dataset and corresponding PCs derived from 100 resampled subsets, each including 80% of participant scans selected without replacement. Component congruence quantifies the similarity between loading vectors, with higher values indicating more reproducible spatial patterns. The green dashed line indicates high congruence (0.95), and the red dashed line indicates the predefined stability threshold (0.85). The first eight PCs had median congruence coefficients above 0.85 and were therefore considered stable and retained for downstream analyses. PCs 9–14 showed lower and more variable congruence, indicating the heterogeneity of the data and less stable spatial patterns across resampled subsets.*

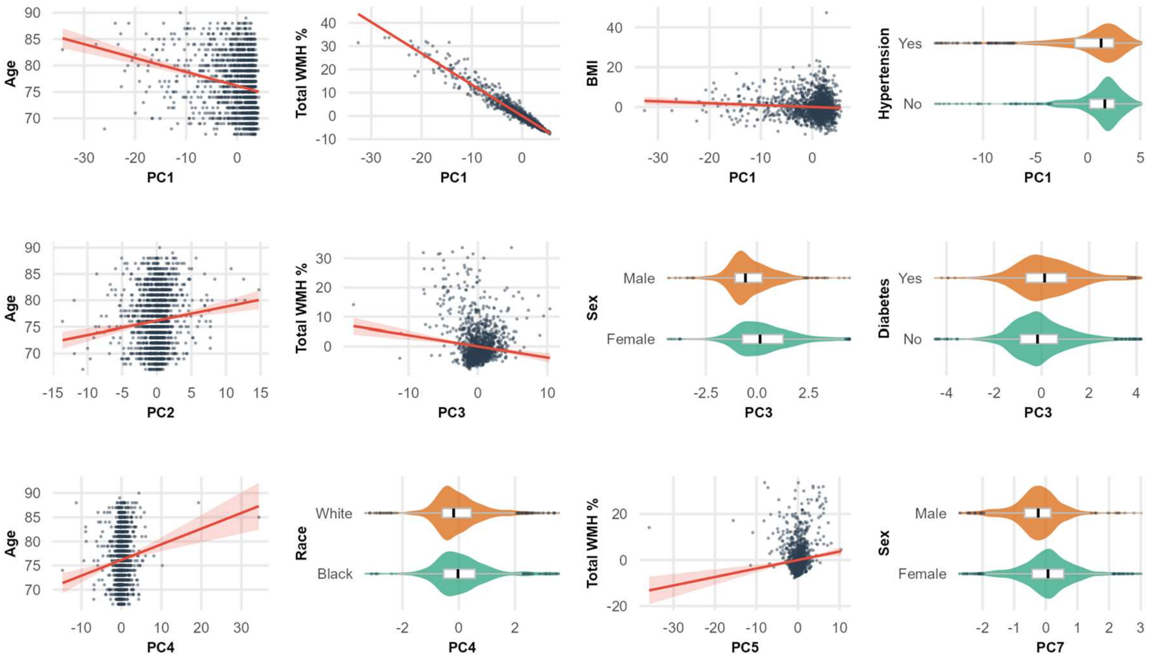


**Supplementary Figure 4**. Relationship of Significant Correlation between Principal Component and Risk Factors

*Scatter plots and violin plots showing significant correlations between white matter hyperintensity (WMH) principal components (PCs) and risk factors. For age and sex, unadjusted associations are presented. For all other variables, associations are adjusted for age and sex using the residualization method, where both variables are regressed on covariates and residuals are plotted. Violin plots display the distribution of PC scores (or residualized PC scores) by category, with embedded box plots showing median, interquartile range, and outliers; x axis trimmed 2.5th–97.5th percentiles of the data for visualization while full data is used for all statistical analysis. Red lines in scatter plots represent linear regression fits with 95% confidence intervals.*

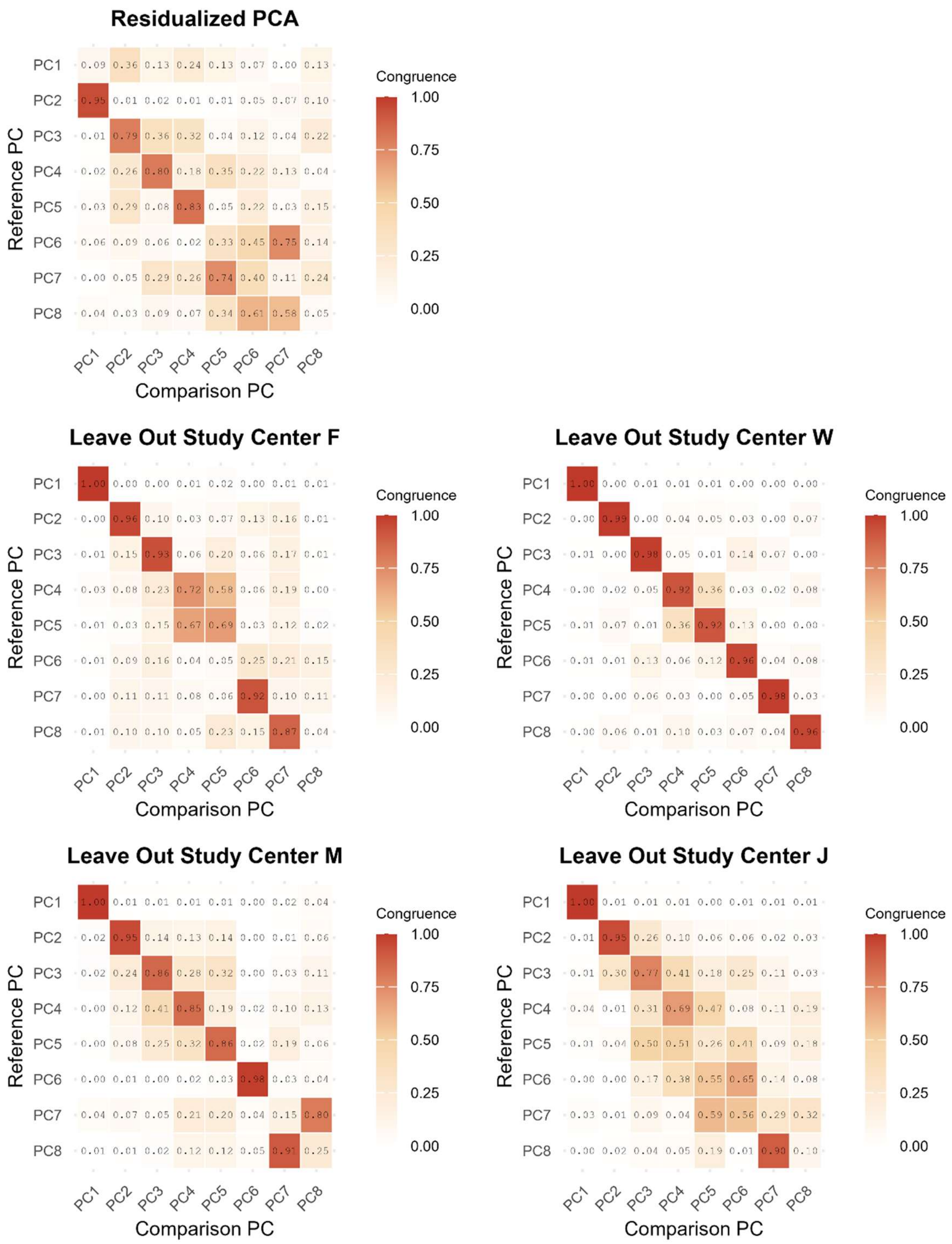


**Supplementary Figure 5**. Tucker’s congruence coefficients between the original PCA solution and alternative decompositions

*Each panel shows the matrix of Tucker's congruence coefficients between the eight retained components of the original PCA solution (reference, y-axis) and the components of an alternative decomposition (comparison, x-axis). Cell values are absolute congruence coefficients, as the sign of a principal component is arbitrary and only magnitude is informative. The color scale denotes |Tucker's φ| ranging from 0 (no correspondence) to 1 (identical loading pattern). Values on the diagonal reflect correspondence between identically numbered components, whereas high off-diagonal values indicate that a component was recovered but reordered in the alternative solution; the best-match congruence reported in the text is the highest coefficient in each reference row after optimal one-to-one assignment of components. (a) Residualized PCA, performed after regressing each regional WMH percentage on total WMH volume. (b–e) Out-of-fold site-wise PCA: each panel corresponds to one ARIC field centers, and shows the solution derived from the remaining three centers, which was used to score the participants of the held-out center.*

**Supplementary Table 2**. Sensitivity Analysis using Inverse Probability of Attrition Weighting: Linear Mixed-Effects Model Estimates for Principal Components with Significant Associations with Cognitive Performance

| Predictor | Global | Executive | Language | Memory |
|---|---|---|---|---|
| PC1 | 0.0132[0.0043, 0.0222](0.0154) | 0.0157[0.0058, 0.0256](0.0144) | 0.0097[-0.0005, 0.0199](0.1896) | 0.0041[-0.0063, 0.0146](0.6094) |
| PC3 | 0.0065[-0.0144, 0.0275](0.6663) | 0.0048[-0.0183, 0.0280](0.7282) | 0.0091[-0.0148, 0.0330](0.5588) | 0.0121[-0.0122, 0.0364](0.6094) |
| PC8 | -0.0043[-0.0291, 0.0204](0.7791) | -0.0215[-0.0489, 0.0058](0.2455) | -0.0127[-0.0410, 0.0156](0.5472) | 0.0143[-0.0145, 0.0430](0.6094) |
| PC1:Time | 0.0025[0.0017, 0.0033](<0.0001) | 0.0025[0.0014, 0.0036](0.0002) | 0.0021[0.0010, 0.0033](0.0028) | 0.0016[0.0005, 0.0027](0.0359) |
| PC3:Time | -0.0041[-0.0061, -0.0020](0.0008) | -0.0009[-0.0037, 0.0018](0.5987) | -0.0055[-0.0084, -0.0026](0.0027) | -0.0040[-0.0066, -0.0013](0.0359) |
| PC8:Time | -0.0043[-0.0067, -0.0018](0.0041) | -0.0026[-0.0059, 0.0007](0.2455) | -0.0028[-0.0063, 0.0007](0.2377) | -0.0012[-0.0045, 0.0020](0.6094) |

*Data shown as coefficient [95% Confidence Interval] (p-value). Models are identical to the primary analysis except that each observation was weighted by the inverse of the participant's estimated probability of having longitudinal follow-up, derived from a logistic model of baseline characteristics. All p-values were corrected for multiple testing across the 8 principal component main effects and 8 principal component × time interaction terms within each cognitive domain using the false discovery rate. PC: principal component.*

**Supplementary Table 3**: Sensitivity Analysis of Linear Mixed-Effects Model Estimates for Principal Components with Significant Association with Cognitive Performance

| | Global | Executive | Language | Memory |
|---|---|---|---|---|
| PC1 | 0.0088[-0.0199, 0.0375](0.6592) | 0.0108[-0.0206, 0.0423](0.6140) | 0.0155[-0.0166, 0.0476](0.6093) | -0.0069[-0.0404, 0.0267](0.8615) |
| PC3 | 0.0026[-0.0150, 0.0201](0.8258) | 0.0043[-0.0152, 0.0238](0.6675) | -0.0002[-0.0202, 0.0198](0.9842) | 0.0084[-0.0123, 0.0290](0.6838) |
| PC8 | -0.0202[-0.0415, 0.0011](0.2030) | -0.0321[-0.0558, -0.0084](0.0420) | -0.0187[-0.0431, 0.0057](0.3568) | 0.0006[-0.0246, 0.0258](0.9630) |
| PC1:Time | 0.0022[0.0014, 0.0031](<0.0001) | 0.0021[0.0010, 0.0033](0.0061) | 0.0020[0.0007, 0.0032](0.0151) | 0.0018[0.0007, 0.0030](0.0291) |
| PC3:Time | -0.0042[-0.0062, -0.0021](0.0007) | -0.0018[-0.0046, 0.0010](0.3857) | -0.0050[-0.0079, -0.0021](0.0131) | -0.0038[-0.0065, -0.0011](0.0516) |
| PC8:Time | -0.0028[-0.0053, -0.0003](0.1242) | -0.0019[-0.0052, 0.0015](0.4346) | -0.0017[-0.0052, 0.0019](0.6093) | -0.0007[-0.0040, 0.0027](0.8615) |

*Data shown in coefficient [95% Confidence Interval] (p-value), all p-values were corrected for multiple testing across the 8 principal component main effects and 8 principal component × time interaction terms within each cognitive domain using the false discovery rate PC: principal component.*

**Supplementary Table 4**. Linear Mixed-Effects Model Estimates Derived from the Residualized Principal Component Analysis

| Predictor | Global | Executive | Language | Memory |
|---|---|---|---|---|
| PC3 | -0.0017[-0.0159, 0.0125](0.9300) | -0.0022[-0.0181, 0.0136](0.7816) | -0.0039[-0.0202, 0.0123](0.8947) | 0.0041[-0.0126, 0.0208](0.8733) |
| PC8 | -0.0136[-0.0333, 0.0060](0.3481) | -0.0116[-0.0333, 0.0101](0.3939) | -0.0239[-0.0463, -0.0015](0.1956) | 0.0069[-0.0161, 0.0300](0.8733) |
| PC3:Time | -0.0028[-0.0046, -0.0011](0.0119) | -0.0014[-0.0037, 0.0010](0.3825) | -0.0046[-0.0070, -0.0021](0.0045) | -0.0026[-0.0049, -0.0003](0.1584) |
| PC8:Time | -0.0024[-0.0048, 0.0001](0.2423) | -0.0041[-0.0074, -0.0009](0.1029) | -0.0005[-0.0040, 0.0030](0.8947) | -0.0016[-0.0048, 0.0017](0.8733) |

*Data shown as coefficient [95% Confidence Interval] (p-value). Principal components were derived after regressing each regional WMH percentage on total WMH volume. The component labels in the Predictor column do not refer to the component numbering of the residualized solution; each component of the residualized solution was matched to the component of the original solution with which it showed the highest Tucker's congruence coefficient (Supplementary Figure 5a), and is reported under that original component's label. Because the sign of a principal component is arbitrary, coefficients were multiplied by the sign of the corresponding congruence coefficient so that the direction of association is directly comparable with the primary analysis; p-values are unaffected by this operation. All p-values were corrected for multiple testing across the 8 principal component main effects and 8 principal component × time interaction terms within each cognitive domain using the false discovery rate. PC: principal component.*

**Supplementary Table 5**. Linear Mixed-Effects Model Estimates Derived from the Out-of-Fold, Site-wise Principal Component Analysis

| Predictor | Global | Executive | Language | Memory |
|---|---|---|---|---|
| PC1 | 0.0148[0.0078, 0.0218](0.0003) | 0.0162[0.0085, 0.0240](0.0007) | 0.0130[0.0051, 0.0210](0.0104) | 0.0053[-0.0030, 0.0135](0.4398) |
| PC3 | 0.0103[-0.0081, 0.0286](0.3955) | 0.0109[-0.0095, 0.0314](0.4933) | 0.0012[-0.0197, 0.0221](0.9777) | 0.0148[-0.0066, 0.0363](0.4398) |
| PC8 | -0.0134[-0.0297, 0.0028](0.2405) | -0.0212[-0.0393, -0.0031](0.1165) | -0.0069[-0.0258, 0.0119](0.7816) | -0.0048[-0.0240, 0.0144](0.6891) |
| PC1:Time | 0.0021[0.0013, 0.0030](0.0000) | 0.0020[0.0009, 0.0032](0.0038) | 0.0019[0.0007, 0.0031](0.0104) | 0.0017[0.0006, 0.0029](0.0276) |
| PC3:Time | -0.0038[-0.0059, -0.0016](0.0031) | -0.0010[-0.0038, 0.0019](0.5499) | -0.0048[-0.0078, -0.0018](0.0104) | -0.0037[-0.0065, -0.0009](0.0529) |
| PC8:Time | -0.0018[-0.0040, 0.0005](0.2555) | -0.0015[-0.0045, 0.0014](0.4933) | -0.0016[-0.0047, 0.0016](0.6732) | 0.0006[-0.0024, 0.0036](0.6891) |

*Data shown as coefficient [95% Confidence Interval] (p-value). For each of the four ARIC field centers in turn, principal components were derived from the remaining three centers and applied to score the participants of the held-out center; the four sets of projected scores were then combined so that no participant was scored using a solution derived in part from their own center. The component labels in the Predictor column do not refer to the component numbering of any individual site-wise solution; within each fold, components were matched to those of the original solution by Tucker's congruence coefficient (Supplementary Figure 5b–e) and sign-aligned before the scores were combined, so that each column corresponds to the original component under which it is labeled. All p-values were corrected for multiple testing across the 8 principal component main effects and 8 principal component × time interaction terms within each cognitive domain using the false discovery rate. PC: principal component.*
